# 3D-printed, Citrate-based Bioresorbable Vascular Scaffolds for Coronary Artery Angioplasty

**DOI:** 10.1101/2023.10.27.564432

**Authors:** Yonghui Ding, Liam Warlick, Mian Chen, Eden Taddese, Caralyn Collins, Rao Fu, Chongwen Duan, Xinlong Wang, Henry Ware, Cheng Sun, Guillermo Ameer

**Affiliations:** Centre for Advanced Regenerative Engineering (CARE), Northwestern University, Evanston, IL 60208, USA; Department of Biomedical Engineering, Northwestern University, Evanston, IL 60208, USA; Department of Mechanical Engineering, Northwestern University, Evanston, IL 60208, USA; Department of Surgery, Feinberg School of Medicine, Northwestern University, Chicago, IL 60611, USA

## Abstract

Fully bioresorbable vascular scaffolds (BVSs) were designed to overcome the limitations of metallic drug-eluting stents (DESs). However, current polymer-based BVSs, such as Abbott’s Absorb, the only US FDA-approved BVS, struggle with increased strut thickness (150 μm for Absorb) and exacerbated tissue inflammation, leading to inferior clinical performance compared to metallic DESs. Here we develop a drug-eluting BVS (DE-BVS) through the innovative use of photopolymerizable, citrate-based materials and high-precision additive manufacturing process. Bare BVS with a clinically relevant strut thickness of 62 μm can be produced in a high-throughput manner, i.e. one BVS per minute. By modulating the coating polymer and structure, we achieve a controlled release of anti-restenosis drug of everolimus from DE-BVSs. We show the mechanical competence of DE-BVS and the successful deployment in swine coronary arteries using a custom-built balloon catheter delivery system. We further demonstrate that BVS and DE-BVS remain safe and effective to keep the vessel patency, induce limited inflammation, and facilitate the recovery of smooth muscle and endothelial tissues over 28 days implantation in swine coronary arteries. All these evaluated pre-clinical performances are largely comparable to the commercial XIENCE™ DES (Abbott Vascular).

## Introduction

Atherosclerotic coronary artery disease (CAD) is responsible for significant morbidity, mortality, and high healthcare costs worldwide. Endovascular therapies, such as the implantation of vascular stents after angioplasty, have become the mainstream treatment for CAD. Bare-metal stents act as a mechanical scaffold to restore blood flow; however, they are associated with high restenosis rates (20-60%)^1, 2^. Drug-eluting stents (DESs) loaded with anti-stenosis drugs, such as sirolimus and everolimus, effectively reduced the restenosis rate to < 5%^3^. However, the permanent presence of metallic frame in the blood vessel causes a range of side effects, including impaired vasomotion, compromised adaptive arterial remodeling, and long-term foreign-body responses^4-6^. Also, when permanent stents fail, the revision surgery to revascularize tissue is complicated and associated with dismal outcomes. These problems associated with permanent metal stents have prompted the development of bioresorbable vascular scaffolds (BVSs). Metal-based BVSs, often made of magnesium or iron-based alloys, have not been clinically adopted due to their rapid resorption or detrimental corrosion products, which leads to frequent post-surgical restenosis^7^. Polymer-based BVSs have emerged as a promising solution for providing initial support to prevent blood vessel recoil, slowly degrading to eliminate residual foreign materials, and eventually restoring blood vessel functions such as vasomotion^8^. However, current polymer-based BVSs, such as Absorb GT1 (Abbott Vascular, Santa Clara, CA), the only U.S. Food and Drug Association (FDA)-approved BVS, are associated with a high incidence of late thrombosis (3.5% vs 0.9% at 2 years), major adverse cardiac events (11% vs 7.9% at 2 years), and require prolonged dual-antiplatelet therapy to prevent clotting when compared to metal DES^9,10^. Probable causes of these problems include: 1) the oxidative stress and exacerbated tissue inflammation induced by the degradation product of poly (l-lactic acid) (PLLA); and 2) the increased strut thickness (150 μm) compared to metallic DES (61-80 μm in struts thickness) leading to larger disturbance of local blood flow. The thickness of the stent struts also limits the number of patients that can be treated with these devices.

We tackle the aforementioned challenges via the implementation of a two-fold strategy. To address the inflammatory responses to PLLA, we have been investigating the use of citrate-based polymers, such as polydiolcitrates to fabricate BVSs. Polydiolcitrates have demonstrated intrinsic antioxidant properties that reduce chronic inflammatory responses associated with PLLA^11^. Of note, polydiolcitrates have been recently used for the fabrication of FDA-cleared biodegradable implantable medical devices currently used in musculoskeletal surgeries^12^. Furthermore, we and others have shown that polydiolcitrates have intrinsic thromboresistant properties and support the formation of a functional endothelium.^13-15^. To address the adverse impact of increased strut thickness, we refined the high-precision additive manufacturing process flow to fabricate drug-eluting BVSs (DE-BVSs) with a clinically relevant strut thickness of less than 100 μm.

The fundamental understanding of material properties and manufacturing as they relate to *in vivo* performance and clinical feasibility of a BVS is crucial. In this work, to maximize clinical feasibility, we first improved our manufacturing efficiency, resulting in the concurrent fabrication of at least 8 BVSs with a strut thickness of 65 μm and a length of 10 mm within 7 minutes, i.e. one scaffold per minute. To ensure that we can deliver anti-restenosis drugs, on par with current DES on the market, we developed a biodegradable citrate-based polymer coating that can be sprayed onto the 3D-printed BVSs to achieve programmable drug loading and controlled release rates. This polymer coating method can be applied to any tissue regeneration porous scaffold to achieve controlled drug release. Additionally, to allow for the successful deployment of the BVSs, we have developed a customized balloon catheter assembly. Finally, we demonstrate the successful deployment, safety, and performance of BVS and drug-eluting BVS (DE-BVS) within the swine coronary arteries. At 28 days, both BVSs and DE-BVSs demonstrated equivalence relative to the clinically used cobalt chromium (CoCr) DES, i.e. XIENCE Pro DES from Abbott Vascular. Therefore, we show that cirate-based BVS is safe and effective and the additive manufacturing fabrication of DE-BVSs with strut dimensions comparable to those of metal DES is feasible.

## Results

### MicroCLIP allows high-throughput printing of BVS with reduced strut dimensions

The poly (1,12-dodecanediol citrate) (PDC) and poly (1,8-octanediol citrate) (POC) pre-polymer was prepared by the condensation reaction between citric acid and 1,12-dodecanediol or 1,8-octanediol, respectively (as shown in **Supporting Information Figure S1A**). The analysis of PDC molecular structure by mass spectrometry confirmed the presence of C_24_H_38_O_14_ (Mw 550), which is the combination of one 1,12-dodecanediol and two citrate acid, and the C_18_H_30_O_7_ (Mw 358) as the repeating unit (**Figure S1B**). The photo polymerizable PDC and POC were prepared by incorporating methacrylate groups into the PDC or POC pre-polymer (**Figure S1A**). The presence of the C=C bonds in the methacrylate PDC (mPDC) and methacrylate POC (mPOC) were validated by the ^1^H Nuclear Magnetic Resonance (NMR) spectrum (**Figure S1C**). The 3D-printable ink was formulated by mixing mPDC pre-polymer (75 wt%) with a photo initiator I 819, a co-initiator EDAB, and ethanol as a dilutant. The additive manufacturing technology, referred to as microscale continuous liquid interface production (MicroCLIP) (**Figure 1A**), developed in this study offers uniformity of the projected UV light intensity with the pixel size of 4.0 µm. These features allow the high-throughput fabrication of 8 BVSs with length of 10 mm per printing within 7 minutes (one stent per minute) (**Figure 1B**) and the high fabrication fidelity of BVSs with struts as thin as 62 µm (**Figure 1C**).

**Figure 1.**
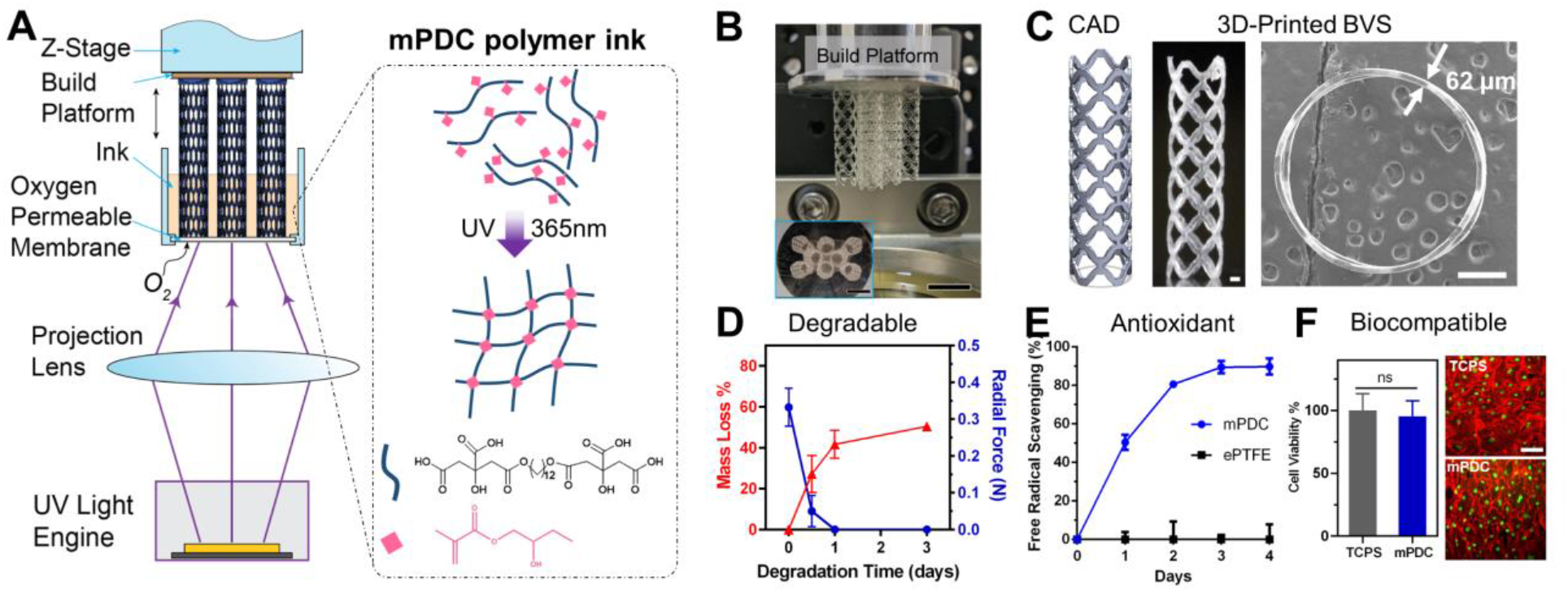
Fabrication and characteristics of the citrate-based, 3D-printed bioresorbable vascular scaffold (BVS). (A) Schematic of micro continuous liquid interface production (MicroCLIP) process for the fabrication of BVS using mPDC polymer ink. (B) Photo shows simultaneous 3D printing of 8 BVSs. (C) BVS CAD model and 3D-printed BVS with 62 ± 3μm strut thickness (left: optical image; right: top view SEM image) Scale bars: 500 μm. (D-F) 3D-printed, citrate polymer-based BVS has biodegradable, antioxidant, and biocompatible properties. (D) Accelerated degradation of BVS, i.e. mass loss (%) and radial force at 30% compression as the functions of degradation time in 0.1 mM NaOH (n = 6). (E) Free radical (i.e. DPPH) scavenging capability of BVS as compared to ePTFE (n = 3). (F) Viability (MTT assay) and cell morphology (red: F-actin, green: nuclei) of HUVECs that were grown on TCPS and mPDC. ns: no significance. n = 3. Scale bar: 20 μm.

The key properties of 3D-printed BVSs, including degradation, antioxidant properties, and biocompatibility, were characterized. The mass loss and the reduction in the radial strength were observed following accelerated degradation in the 0.1 mM sodium hydroxide (**Figure 1D**). The 3D printed BVSs retained the intrinsic antioxidant properties of PDC^11^, as shown by their strong capabilities to scavenge the free radicals (**Figure 1G**). In contrast, limited free radical scavenging was associated with medical grade expanded polytetrafluoroethylene (ePTFE), a commonly used materials for vascular grafts and stent grafts. The biocompatibility with vascular endothelial cells was validated by the high viability and healthy cell morphology when cells were grown on the 3D-printed mPDC, which were comparable to the cells grown on the tissue culture polystyrene (TCPS) (**Figure 1F**).

### mPOC coating of BVS enables controlled release of everolimus

DES consisting of a metal stent with a thin polymer coating as the carrier of anti-stenosis drugs has been developed to combat intimal hyperplasia. Current approaches to control the release of a drug from a stent have challenges that impact endothelial tissue recovery. Here, we applied mPOC as the drug carrying polymer coating onto 3D-printed BVSs and show that the controlled release of everolimus was achieved (**Figure 2**). The drug-eluting BVSs (DE-BVSs) were fabricated by sequentially depositing a polymer-drug layer (mPOC with everolimus) and a barrier layer (mPOC only) through an ultrasonic spray coating method (**Figure 2A**). A conformal and uniform coating with a smooth surface finish was obtained on both luminal and abluminal surfaces of the DE-BVSs as shown by scanning electron microscopy (SEM) images (**Supporting Information Figure S2)**. The strut thickness was increased from 62 ± 3 μm for the bare BVS to 99 ± 4 μm for the DE-BVS (drug layer and barrier layer). The successful incorporation of everolimus within the coating of drug layer was confirmed by ATR-FTIR analysis (**Supporting Information Figure S3**). The characteristic peaks for pure everolimus, including 1643 cm^-1^ (C=O), 1451 cm^-1^ (C-H), and 990 cm^-1^ (C-H)^16^, were detected at the spectra of DE-BVS (drug and drug+barrier) while they were not visible at the spectra of BVS with pure polymer coating not containing the drug.

**Figure 2.**
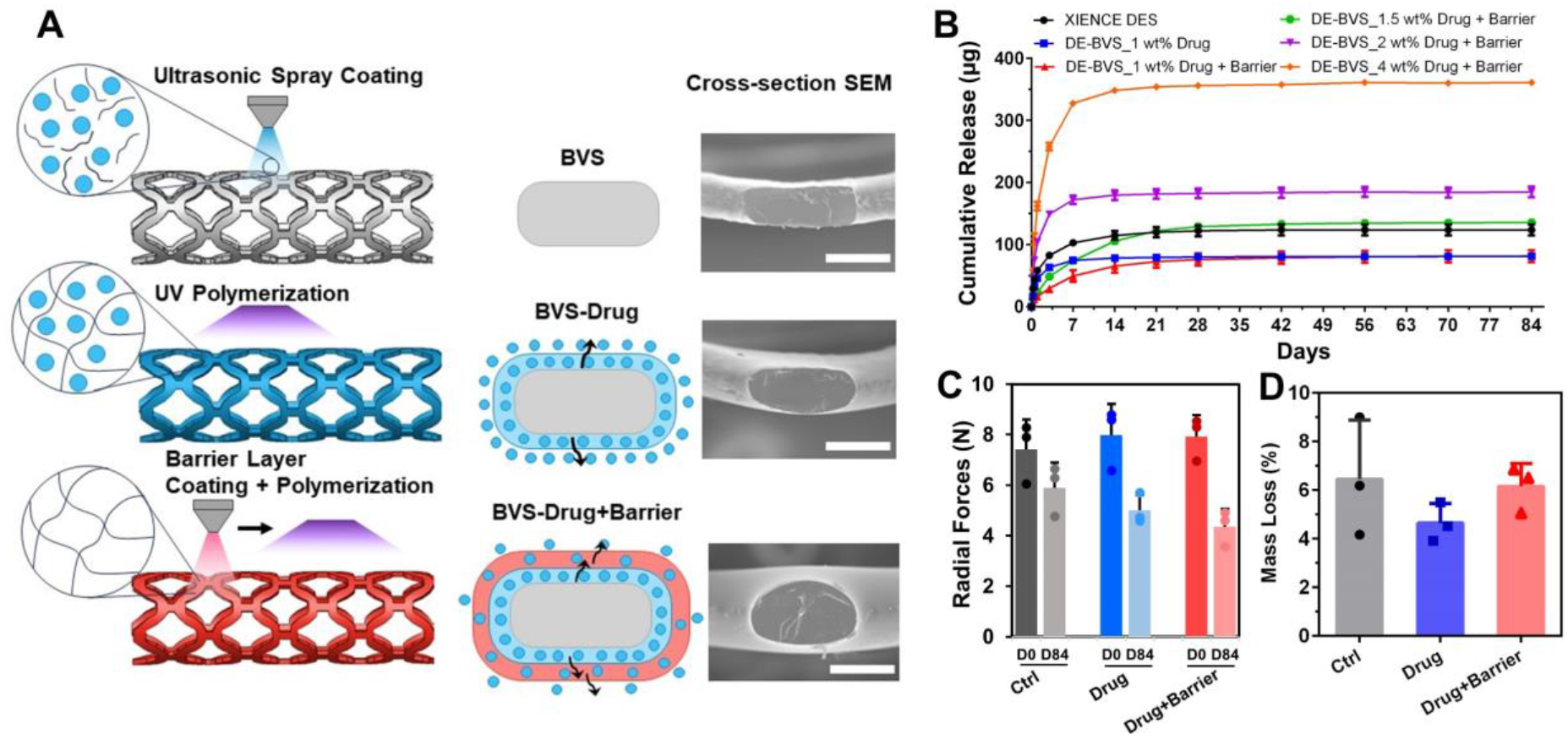
Fabrication of drug eluting BVS (DE-BVS) with strut thickness under 100 μm and programmable drug eluting profiles. (A) Scanning electron microscope (SEM) cross-sectional views of radial strut thickness of bare BVS (62 ± 3 μm) and DE-BVS with coatings of drug (77 ± 4 μm), and DE-BVS with coatings of drug+barrier (99 ± 4 μm). Scale bar: 100 μm. (B) Various drug eluting profiles over 84 days were achieved by manipulating coating structures and loading concentrations (1, 1.5, 2, and 4 wt%) of drugs (n = 4, *p* > 0.05). (C, D) Degradation behaviors in regards of (C) radial forces at 50% compression at day 0 (D0) and day 84 (D84) as well as (D) the mass loss of DE-BVS in the PBS at 37°C over 84 days (n=3, *p* > 0.05).

Under normal physiological conditions, a set of release profiles were achieved with DE-BVSs by programming the order of the coatings (drug *versus* drug+barrier) and the loading concentrations of drug (**Figure 2B**). For example, a burst release was observed for the DE-BVS with drug layer only, with 92% of everolimus being released within 7 days. A more sustained release was achieved for the DE-BVS with drug + barrier coatings, with only 60% and 80% of everolimus being released within 7 and 14 days, respectively. Notably, the commercial XIENCE DES showed the release of 83% and 93% of everolimus within 7 and 14 days. In addition, the loading concentration of everolimus, such as 1, 1.5, 2, and 4 wt%, into the coating solution could be tuned to control the total amount of everolimus being loaded onto and released from the DE-BVS. To closely match the total amount of everolimus released from commercial XIENCE DES, the DE-BVS with the loading concentration of 1.5 wt% was selected for evaluation *in vitro* and *in vivo*.

Following the drug release in PBS with 10% ethanol, pH 7.4, 37°C for 84 days, the degradation of DE-BVSs was assessed by measuring the changes of their radial forces and mass. It was found that the radial force per 8 mm long BVS dropped by 20 - 45% within 84 days (**Figure 2C**), accompanied by a mass loss of 4.6 - 6.5% (**Figure 2D**).

### DE-BVS inhibits proliferation of smooth muscle cells (SMCs) with limited impact on endothelial cells (ECs)

The polymer coating of current DES and the released everolimus interact with surrounding vascular tissues by inhibiting the proliferation of SMCs to reduce the restenosis rate while delaying the recovery of ECs and subsequent vascular healing^17^. The vascular cell responses were evaluated by seeding and growing human umbilical vein endothelial cells (HUVECs) and human aortic smooth muscle cells (HAoSMCs) on 3D-printed mPDC substrates with three types of coatings: 1) mPOC polymer without drug (control), 2) mPOC polymer coating with 1.5 wt% everolimus, and 3) mPOC polymer coating with 1.5% everolimus plus a mPOC barrier layer (drug + barrier) (**Figure 3**). The high cell viability and healthy cell morphologies of HAoSMCs and HUVECs were observed on the control substrate, suggesting the good biocompatibility of mPOC polymer as a coating for scaffolds. The released everolimus from the polymer/drug coating prominently inhibited the proliferation and viability of both HAoSMCs and HUVECs. Interestingly, the coatings of drug + barrier significantly inhibited SMC viability and spreading, while it showed limited impact on EC viability and morphology. These *in vitro* results suggested the importance of controlled release of everolimus from DE-BVSs in modulating the selective responses of SMCs and ECs.

**Figure 3.**
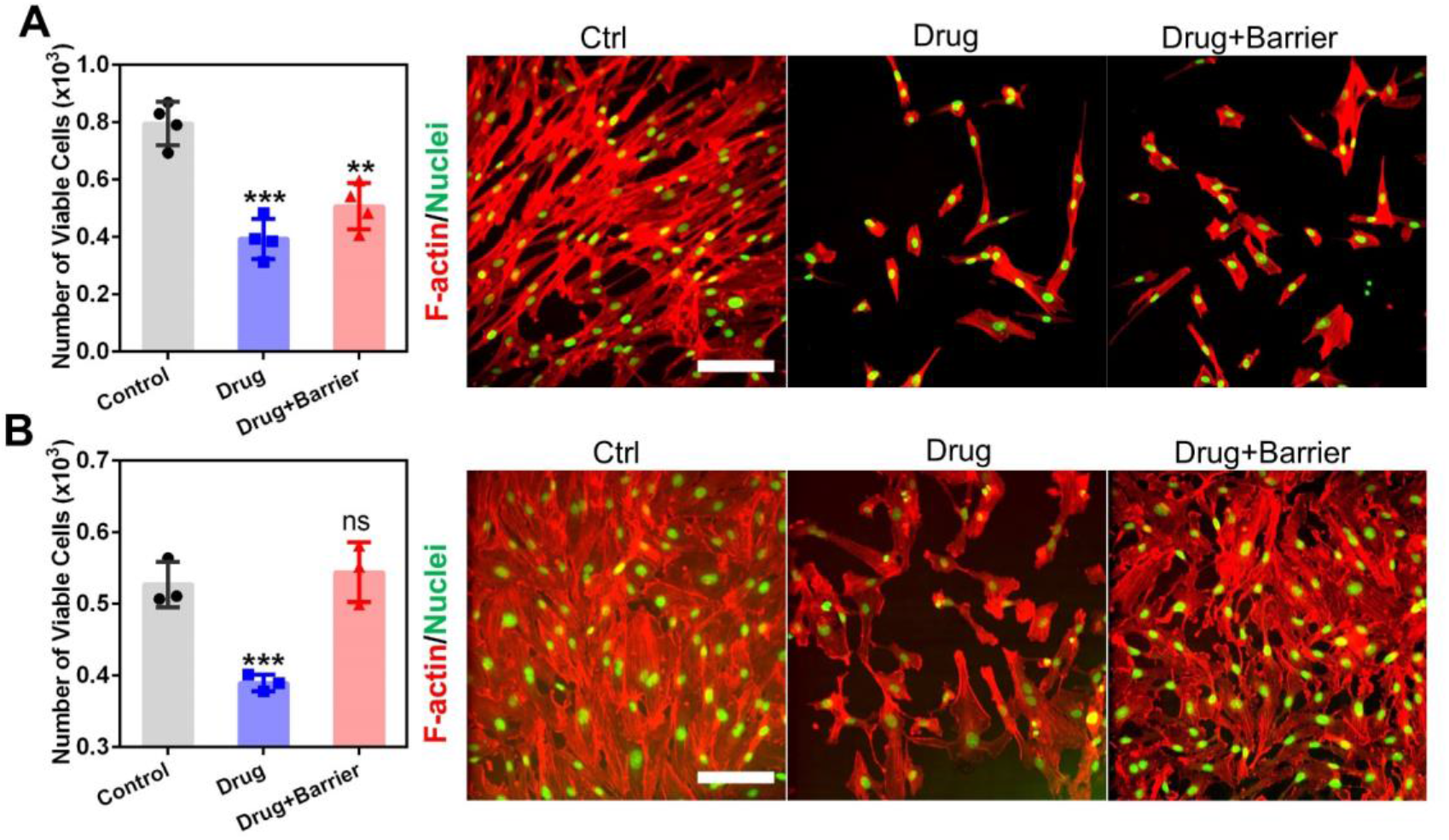
*In vitro* responses of (A) human aortic smooth muscle cells (HAoSMCs) and human umbilical vein endothelial cells (HUVECs) to DE-BVSs. Cell viability was evaluated by Alamar Blue Assay (n = 3 or 4; ** *p* < 0.01 and *** *p* < 0.001 vs. control). Cell morphologies were observed by confocal microscope following the immunostaining of F-actin and nuclei. Scale bars: 200 μm.

### A customized balloon catheter system enables the *in vivo* deployment of BVSs

To prepare for the *in vivo* evaluation of scaffolds, we investigated the possibility of deploying our 3D-printed BVSs and DE-BVSs using the standard catheterization procedures used in current clinical practice of interventional cardiology. We first optimized the scaffold geometry, and conditions of 3D printing and post-processing to ensure the mechanical competence of scaffolds. As a result, 3D-printed scaffolds could withstand cycling mechanical loading, including radial compression and expansion between the original diameter of 3.0 mm and final, crimped diameter of 1.1 mm without strut fracture (**Figure 4A, B and Supporting Information Figure S4**). Interestingly, we observed spontaneous self-expansion and an increase of scaffold diameter following the scaffold compression onto the balloon catheter (**Supporting Information Figure S4)**. This self-expansion can potentially lead to the dislodgement between the compressed scaffold and the balloon during the advancement of the balloon catheter toward the targeted location inside the artery.

**Figure 4.**
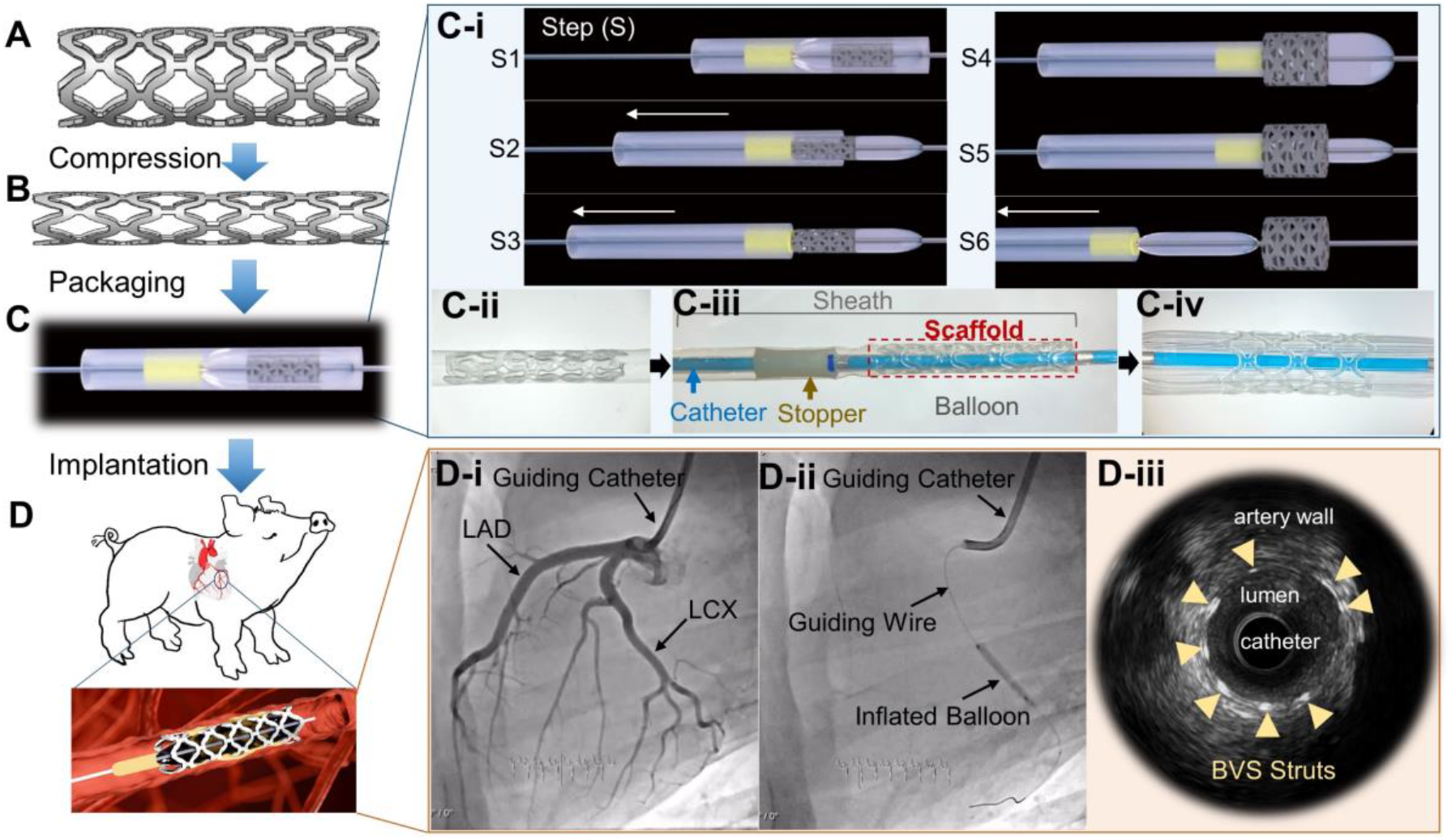
The custom-built BVS delivery system enables the deployment of BVS into the swine coronary artery. The fabricated scaffolds (A) were compressed (B), assembled onto the custom-built delivery system (C), and implanted into the swine coronary artery (D). (C-i) The schematic illustration of scaffold deployment steps, including step 1 (S1), the assembled scaffold on balloon catheter; S2, the retraction of the sheath with scaffold stopped by the stopper; S3, the release of the scaffold from the sheath; S4, the inflation of the balloon along with the expansion of the scaffold; S5, the deflation of the balloon; and S6, the retraction of the balloon catheter. (C-ii) Digital image shows the compressed scaffold locked inside the catheter sheath. (C-iii) Digital image shows the scaffold assembled onto the custom-built delivery system. (C-iv) Digital image shows the expanded scaffold on the balloon catheter. (D-i) X-ray images from angiography show the left anterior descending coronary artery (LAD) and left circumflex artery (LCX) of a swine. (D-ii) X-ray image shows the inflated balloon to expand the scaffold in the LCX. (D-iii) The intravascular ultrasound (IVUS) image shows scaffold struts (as indicated by arrows) within the coronary artery following implantation.

To address this issue, a customized balloon catheter system was developed. A modification of a commercial balloon angioplasty catheter was achieved by introducing two components: the locking sheath and a stopper (**Figure 4C**). The locking sheath, made by medical-grade expanded poly (tetrafluoroethylene), was used to keep the compressed scaffold in place (**Figure 4C-ii**). The locked scaffold within the sheath was assembled on the balloon catheter with a stopper at the far end of the balloon (**Figure 4C-iii**). During the deployment process (**Figure 4C-i**), the assembled device was first advanced into the targeted location inside the artery (Step 1 or S1). While retracting the sheath, the scaffold was first moved along with the sheath until it was stopped by the stopper, which was made by a thin tube of medical grade polyetheretherketone (S2), leading to the release of the scaffold from the locking sheath (S3). The scaffold was then expanded against the surrounding vessel wall following the inflation (S4) of the balloon. Finally, following the deflation (S5) of the balloon, the entire customized balloon catheter system was retracted and withdrawn from the body (S6).

Using this customized balloon catheter system and procedure, 3D-printed scaffolds, including 8 BVSs and 8 DE-BVSs, as well as 8 commercial everolimus-eluting DES (Abbott’s XIENCE™ Pro S) (standard of care reference devices) were successfully deployed into coronary arteries of 8 domestic healthy swine (**Figure 4D**). Each wine received three implants (random combinations of BVS, DE-BVS and XIENCE stent) into in three main coronary arteries, including left anterior descending coronary artery (LAD), left circumflex artery (LCX), and right coronary artery (RCA). The angiographic images revealed the targeted arteries of LAD and LCX of a swine (**Figure 4D-i**). A BVS was successfully expanded and deployed in the LCX of a swine (**Figure 4 D-ii**). Although the BVS was not visible under fluoroscopic imaging due to the intrinsic radiolucency of most polymer materials, BVS struts showed strong contrast under intravascular ultrasound (IVUS) (**Figure 4D-iii**), which allows us to tack the scaffold *in vivo*. There were no complications such as acute thrombosis, dissection, malposition, or collapse during the procedures identified by immediate angiography and IVUS.

### BVSs and DE-BVSs demonstrated comparable efficacy and safety to commercial XIENCE DES in swine coronary arteries

All animals remained healthy throughout the 28-day duration of the study. None of coronary arteries with scaffolds or stents showed evidence of luminal thrombosis on day 28 post-implantation. There was no evidence of myocardial infarction, epicardial hemorrhage, or other obvious abnormalities along the coronary arteries according to gross examination of the heart.

Both BVSs and DE-BVSs induced similar levels of arterial narrowing over 28 days to that induced by XIENCE DES as measured by IVUS (**Figure 5A**). The Movat’s pentachrome and haematoxylin and eosin (H&E) stained tissue sections showed that: 1) the lumen remained patent without any evidence of thrombosis; and 2) all scaffold/stent struts were fully covered by neo-intima tissues with well-organized structures (**Figure 5B**). The quantitative analyses showed that there were no statistically significant differences in luminal area, neointimal area, neointimal thickness, and area stenosis when comparing BVS or DE-BVS with the XIENCE DES (**Figure 5C** and **Supporting Information Table S1**). The mean values of neointimal area and area restenosis induced by the XIENCE DES were lower than these induced by BVS and DE-BVS, which was mainly ascribed by the significant arterial remodeling responding to the implantation occurred in 2 out of 8 swine. It is also interesting to find that the neointima area of DE-BVS and XIENCE DES groups were 4.6 ± 4.4 and 2.6 ± 1.2 mm^2^, which were not significantly different from the BVS group, i.e. 3.8 ± 2.2 mm^2^ and *p* > 0.46. Similarly, there was no statistically significant difference in the area restenosis among these three groups, i.e. BVS (62.0 ± 23.8 %), DE-BVS (61.8 ± 17.5 %), and XIENCE DES (48.6 ± 22.1 %); *p* > 0.49. These results suggest that vascular responses to BVS and DE-BVS developed in this study are largely comparable to commercial XIENCE DES at all key measures after implantation for 28 days. Interestingly, the bare polymer BVS displayed comparable levels of restenosis to the DE-BVS with the release of everolimus.

**Figure 5.**
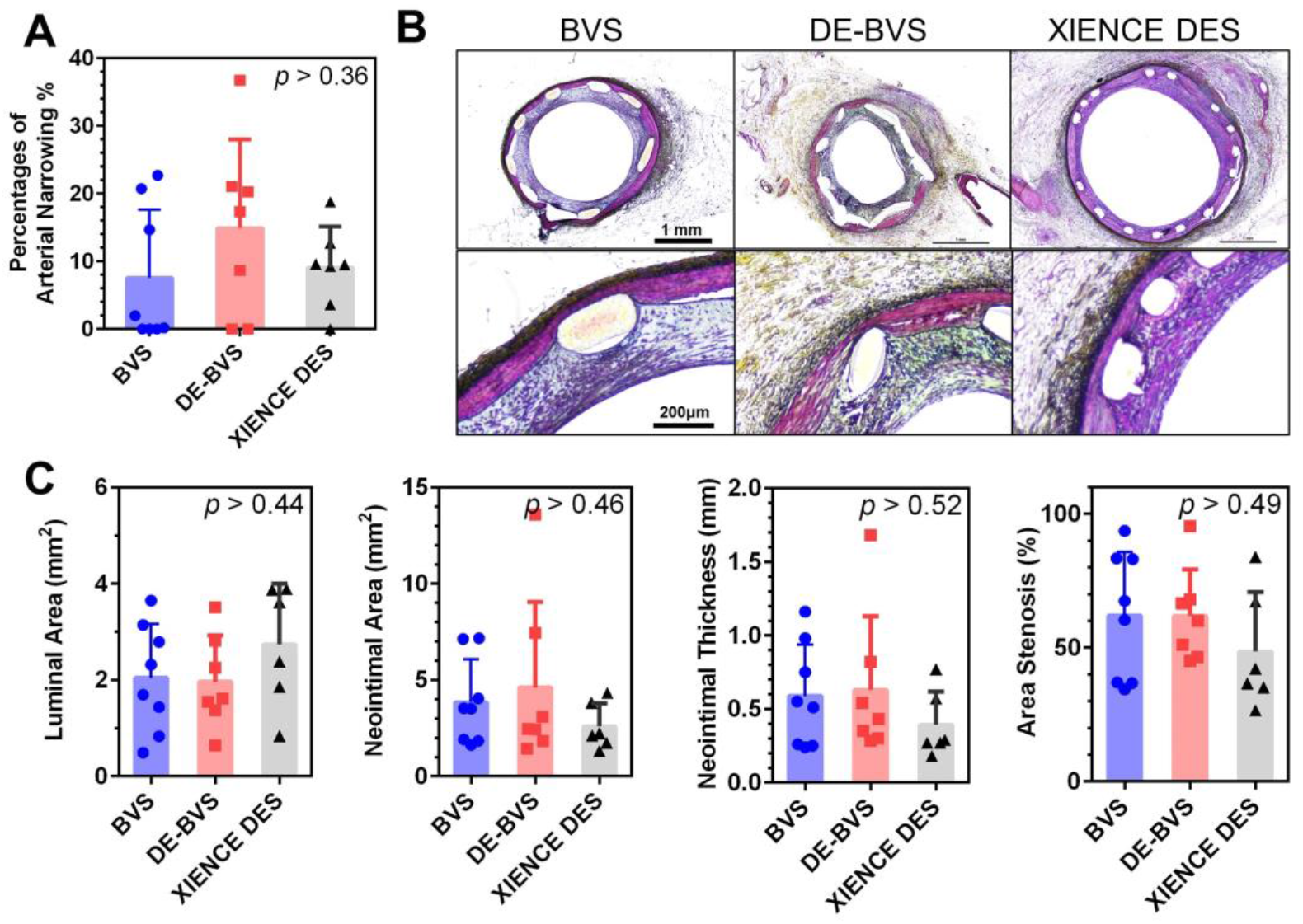
BVSs and DE-BVSs are safe and effective in keeping the arteries patent after implantation for 28 days in swine coronary arteries. (A) Quantification of the arterial narrowing over 28 days as measured via intravascular ultrasound within three stents, i.e. BVS, drug-eluting BVS (DE-BVS), and XIENCE DES. (B) Movat pentachrome stains of swine coronary arteries with scaffolds/stents. (C) Quantification of neointimal thickness, neointimal area, medial area, neointimal stenosis, and inflammation score of stented arteries via the histomorphometry analysis of tissue sections. (n = 8 for BVS; n = 7 for DE-BVS; n = 6 for XIENCE DES.)

### BVSs and DE-BVSs did not induce significant inflammatory responses relative to commercial XIENCE DES

The inflammatory response plays an important role in the healing of the injured arteries and the development of in-stent restenosis. The inflammation was acute and was present only in stented arterial sections. Macrophages are one of the major inflammatory cells that infiltrate into the arterial wall with scaffolds. Therefore, we performed the immunostaining of CD86, a marker of macrophage with a pro-inflammatory (or M1) phenotype, and CD163, a marker of macrophage with a pro-healing (or M2) phenotype (**Figure 6A**). It was found that the infiltrated macrophages were predominantly present in the peri-strut regions. The CD86^+^ and CD163^+^ macrophages largely overlapped with each other, which is in line with the high complexity of macrophage phenotypes^18^. The quantification of the area covered by the CD86^+^ or CD163^+^ macrophages and the ratio of CD163^+^/CD86^+^ suggested that there was no significant difference in the number of macrophages and their phenotypes among three groups, i.e. BVS, DE-BVS and XIENCE DES (**Figure 6B**), which was in consistent with the quantified inflammation score from the histopathological analyses.

**Figure 6.**
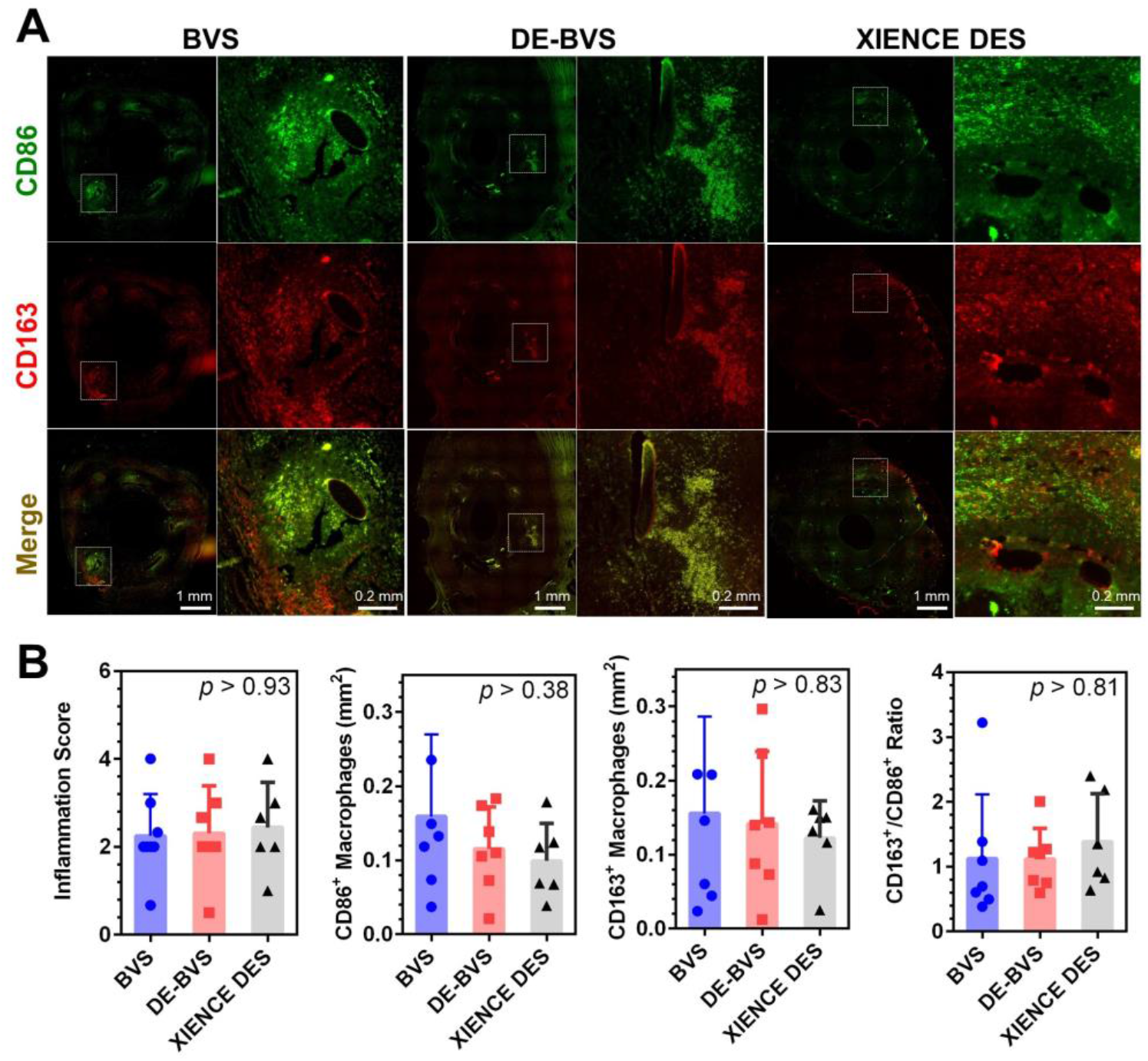
BVSs and DE-BVSs did not induce significant inflammatory responses relative to commercial XIENCE DES. (A) Representative fluorescent images of CD86 and CD163-stained artery sections show the presence of macrophages at peri-strut regions. (B) Quantification of inflammation score (from the H&E-stained artery sections as shown in the supporting information Figure S6), the area coverage of CD86^+^ and CD163^+^-stained macrophages, and the ratio between CD86^+^ and CD163^+^-stained macrophages.

### BVSs and DE-BVSs induced regeneration of smooth muscle and endothelium

To better understand the healing process, we studied the influences of scaffolds/stents on the growth of smooth muscle tissue and re-coverage of endothelium in swine coronary arteries. As shown in **Figure 7A**, the expression of α-SMA was uniform and smooth muscle cells with spindle shape were densely packed in the neointima of all scaffolds/stents. The regenerated smooth muscle tissues demonstrated similar shape and organization to those in the native media layer. The VE-cadherin was used to stain functional endothelial cells. Of note, a thin layer of VE-cadherin^+^ endothelial cells (or endothelium) was observed on the inner most layer of the neointimal tissues for all three groups of scaffolds/stents (as indicated by the white arrow heads on **Figure 7A**). Interestingly, BVSs induced a slightly higher percentage of endothelium coverage (72.4 ± 10.0 % for BVS) than the DE-BVS (60.8 ± 10.0 %) although there was no statistically significant difference (*p* = 0.18). These results suggest that the bare BVS may promote favorable formation of a healthy neointimal layer.

**Figure 7.**
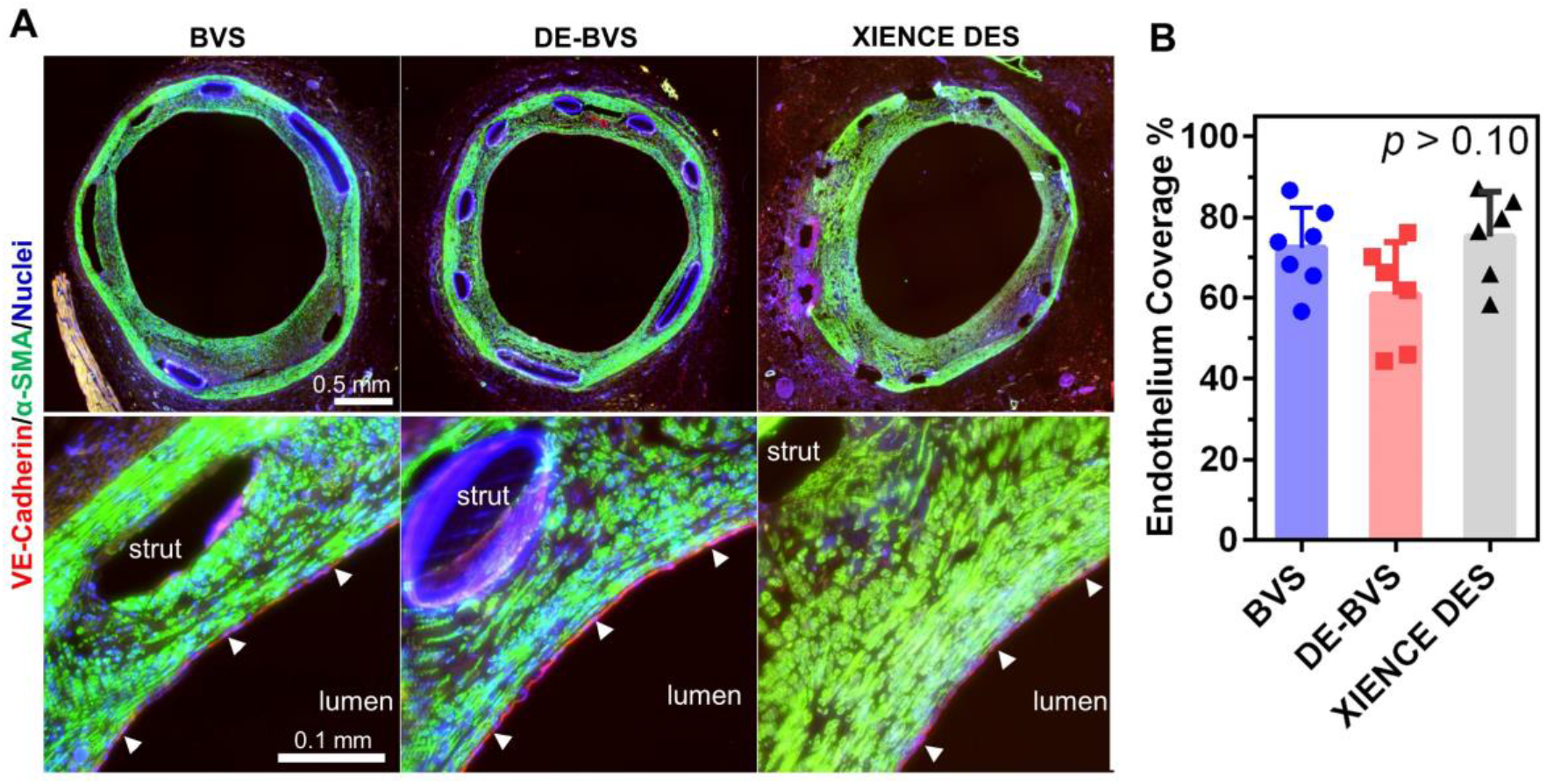
BVSs and DE-BVSs induced the regeneration of smooth muscle and re-coverage of endothelium. (A) Representative fluorescent images of scaffolded/stented arteries by immunostaining of VE-Cadherin (red, marker of endothelial cells), α-SMA (green, marker of smooth muscle cells), and nuclei (blue). The bottom images show enlarged view of selected region from the top images. The regenerated endothelium (the inner most layer of arterial wall) is indicated by the white arrowheads. (B) Quantification of endothelium coverage in scaffolded/stented arteries by measuring and calculating the percentage of region with positive staining of VE-cadherin over the perimeter of lumen.

## Discussion

BVSs were designed to overcome the limitations of permanent metallic DESs, which prevent normal vasomotion, hamper further treatment options in stented segments, and can provoke long-term foreign-body responses^9^. Although multiple scaffolds have been or are in development, the clinical use of these scaffolds, such as the only U.S. FDA-approved Absorb BVS (Abbot Vascular), never gained enough momentum to justify the costs of production and thereby the Absorb BVS was withdrawn from the market in 2017. The inferior clinical performance of current BVS mainly stems from the intrinsic properties of materials used to make BVS, such as PLLA. The unfavorable materials properties include low mechanical strength, requiring larger strut prolife than metallic stents, and the generation of acidic degradation products, leading to a prolonged inflammatory reaction. In addition, current scaffold/stent fabrication technologies, such as laser machining, to fabricate BVS are less effective (cost and time) and lacking the capability of producing patient-specific scaffolds. In this study, we leveraged a citrate-based polymer and a unique 3D printing technology to produce BVSs with biocompatibility, degradation, antioxidant, and drug-eluting properties. The 3D printing technology offers a highly efficient and tailored production of BVSs featuring low profile (sub-100 µm strut thickness and width) and high mechanical strength. The produced BVSs were evaluated in the coronary arteries of the swine model and demonstrated the comparable *in vivo* performance to the commercially available XIENCE DES (Abbot Vascular) at 28 days post-implantation.

The release of anti-restenosis drugs, including parameters such as dosage and release rate, plays a critical role in scaffold/stent performance. Achieving the optimal release profile to prevent the over-proliferation of smooth muscle tissue without retarding the endothelial recovery remains a challenge. The coating polymers used as the drug carrier have large influences on the drug release profile and vascular responses^17, 19-22^. Permanent polymers used in the first generation of DES often lead to late in-stent thrombosis and delayed recovery of endothelium^22^. Despite the promise of biodegradable polymer coatings, the most popular polymer coatings, i.e. polyesters, can cause chronic inflammatory and hypersensitivity reactions because of the generation of acidic degradation products^17^. In this study, for the first time, we report the use of a biodegradable citrate-based polymer as coatings to the BVS. Coatings may consist of the polymer and drug (mPOC/everolimus layer) and/or a second barrier coating consists of just the polymer (mPOC). The result is controlled release of everolimus from the device. Previous work showed that the application of a PLGA blank layer might slow down the drug release too much to combat restenosis, e.g. only 10% of drug released in 30 days^23^. In this study, DE-BVS still released 60% of its everolimus within the first 7 days, which is important to ensure an effective inhibition on SMC proliferation in the early stage following the implantation. Moreover, the degradation product from both the base BVS and the mPOC coating did not trigger significant inflammatory responses at 28 days following the implantation in the swine coronary arteries, an adverse effect often associated with polyester coatings. These findings suggest that the mPOC can serve as a promising coating material for drug-eluting scaffolds/stents, offering good biocompatibility and programmable drug release profiles. In addition, the dosage of the everolimus is another factor to balance the inhibition on SMC and adverse effects on endothelial recovery. In this study, 1.5 wt% loading concentration of everolimus was chosen in order to closely match the total amount of everolimus released from the XIENCE DES (Figure 2B). In the future, measurements of everolimus concentration in arterial tissue and in blood by pharmacokinetic evaluation are needed for a better understanding of the relationship between drug release and vascular tissue response.

The 3D-printed BVS is biocompatible and safe for implantation in swine coronary arteries. Delayed and poor strut coverage by neointimal tissue has been recognized as the primary cause for late and very late stent thrombosis following the placement of metallic DES^4^. This study showed that all BVSs and DE-BVSs were fully covered by neointimal tissue without evidence of thrombosis on day 28 post-implantation. Moreover, there were no significant differences in luminal area, neointimal thickness, and area stenosis among the BVS, DE-BVS and XIENCE DES devices as confirmed by ultrasound imaging examination and histomorphometric analyses. Although the Absorb BVS was not included in this study, previous studies showed that the Absorb BVS induced greater levels of neointimal thickness and stenosis than XIENCE DES in the swine coronary artery at 1 month, e.g. 27.9% stenosis of Absorb vs. 13.5% stenosis of XIENCE DES with *p* = 0.02^24^. These findings suggest potentially improved vascular healing induced by our BVS than Absorb BVS.

Vascular inflammation plays critical roles in scaffolding/stenting outcomes because greater inflammation has been correlated with greater neointimal growth and restenosis^25, 26^. Previous studies have showed that the inflammation induced by Absorb BVS was greater than that induced by XIENCE DES from 6 to 36 months in the swine model, which might be ascribed to the acidic degradation product of Absorb BVS^24^. In this study, the inflammation scores were largely comparable between BVS, DE-BVS and XIENCE DES based on the histomorphometric analyses. Evidence demonstrated the importance of macrophages in vascular healing following scaffolding/stenting. For instance, it was established that M1 macrophages enhance the secretion of pro-inflammatory factors and SMC migration, while M2 macrophages encourage endothelial recovery^27, 28^. In this study, the similar number of CD86^+^ (M1 phenotype marker) and CD163^+^ (M2 phenotype marker) macrophages were observed for all scaffolds/stents suggesting the comparable inflammatory responses.

Endothelium regeneration is an important step toward vascular healing. Given the vital roles played by endothelium in vascular biology^29^, it has been widely recognized that the rapid and functional regeneration of endothelium is critical for achieving long-term patency and biocompatibility of vascular scaffolds/stents^30-32^. In this study, immunohistochemical assessments indicated that the 70-80% coverage of endothelium occurs on day 28 post-implantation for almost all scaffolded/stented arteries. Previous studies have shown poor endothelium recovery with the Absorb BVS than with metallic DES in a rabbit model^33^. Despite the significant efforts made to facilitate endothelium adhesion, proliferation, and migration through a variety of strategies^31, 32^, the mechanism of endothelium regeneration on vascular scaffolds/stent is still unclear^30, 34^. A recent study found that blood circulating monocytes would adhere to the luminal surfaces of acellular vascular graft, differentiate into a mixed endothelial (CD144^+^) and macrophage (CD163^+^) phenotype, and further develop into mature endothelial cells *in vitro* and in the ovine model^35^. Interestingly, we had similar findings that the regenerated endothelial layers in the scaffolded/stented arteries in 3 out of 8 swine co-expressed the endothelial specific marker of CD144^+^ and the macrophage specific makers CD163^+^/CD86^+^ (**Supporting Information Figure S6**), while regenerated endothelial layers in the other 5 out of 8 swine only expressed the endothelial specific marker of CD144^+^, which might be resulted from the different stages of endothelial differentiation from monocytes. This observation suggests the important roles that inflammation plays in the processes of vascular healing after the placement of scaffold/stent, which deserves more investigation in the future.

In summary, we have designed, fabricated, and tested a biocompatible and regenerative BVS that could release anti-restenosis everolimus in a clinically relevant time frame and dosage. We harnessed a high-resolution and high-speed additive manufacturing process to both produce multiple BVS in parallel, drastically reducing fabrication time for this product, and to reliably print low-profile BVSs with strut thicknesses as small as 62 μm. In doing so, we were able to develop a promising solution for the clinically relevant production of patient specific scaffolds. Additionally, we were able to produce these scaffolds and their coatings out of a citrate-based biodegradable polymer. By utilizing mPOC coatings as both drug carrier and barrier layers, a controlled release profile of everolimus can be achieved with 3D-printed scaffolds. Beyond showing favorable biocompatibility and intrinsic anti-oxidant properties, these scaffolds were deployed successfully in swine coronary arteries with the use of a custom-built deployment system and process. These scaffolds demonstrated safety and efficacy to maintain the vessel patency for 28 days after implantation. Moreover, the BVSs and DE-BVSs tested here favored vascular healing processes by promoting rapid strut coverage by neointimal tissues and endothelium recovery while limiting the degree of stenosis. Overall, the BVSs and DE-BVSs developed here demonstrate largely comparable pre-clinical performance in swine coronary arteries as the commercial XIENCE DES, which is the most widely used coronary stent in clinic. In the future, a long-term follow-up (≥ 6 months) and evaluation in atheromatous animal models will be performed to gain more translational perspectives. For instance, the change of lumen area following the partial or full resorption of the scaffold could be an important characteristic to track and distinguish it from metallic stents. In addition, the evaluations of pharmacokinetics and mPDC and mPOC polymer degradation *in vivo* are needed to gain better insights into: 1) the drug release profiles in the arterial tissues and blood flow; and 2) the resorption characteristics of scaffolds and the impact on their mechanical function and vascular healing.

## Materials and Methods

### Materials and Devices

All chemicals were obtained from Millipore Sigma unless otherwise noted. Everolimus was purchased from Selleckchem. The control device, XIENCE™ Pro S everolimus-eluting DES was obtained from Abbott Vascular.

#### mPDC and mPOC synthesis and characterization

Methacrylate poly(1,12-dodecamethylene citrate) (mPDC) and Methacrylate poly(1,8-octandiol citrate) (mPOC) was synthesized by following our previously reported protocol^14^. Briefly, citric acid and 1,12-dodecanediol at molar ratio of 2:3 or citric acid and 1,8-octandiol at molar ratio of 1:1 were melted (165 °C, 22 min), co-polymerized (140 °C, 60 min), purified, and freeze-dried to yield PDC of POC pre-polymer. Every 22 g PDC or POC pre-polymer was dissolved in tetrahydrofuran (180 mL) with imidazole (816 mg) and glycidyl methacrylate (17.8 mL), reacted (60 °C, 6 h), purified, and freeze-dried to yield mPDC or mPOC pre-polymer. Then 5 mg mPDC or mPOC pre-polymer was dissolved in 1 ml deuterated dimethyl sulfoxide (DMSO-d6) and characterized using proton nuclear magnetic resonance (^1^H-NMR).

#### Fabrication of BVS

To formulate mPDC ink for 3D printing of BVS, 75 wt.% mPDC pre-polymer was mixed with 2.2 wt.% Irgacure 819, acting as a primary photoinitiator, and 3.0 wt.% Ethyl 4-dimethylamino benzoate (EDAB), acting as a co-photoinitiator, in a solvent of pure ethanol. 3D printing was performed using a homemade micro-continuous liquid production process (MicroCLIP)-based printer. An oxygen-permeable window made of Teflon AF-2400 (Biogeneral Inc., San Diego, CA) was attached to the bottom of the resin vat. A digital micromirror device (DMD, Texas Instruments Inc., Plano, TX) was utilized as the dynamic mask generator to pattern the UV light (365 nm). Projection optics of the printer were optimized to have a pixel resolution of 7.1 µm x 7.1 µm at the focal plane. CAD files of the print part were designed in SolidWorks (Dassault Systèmes, Waltham, MA). The resulting STL files were sliced for a 2D file output by MATLAB (MathWorks Inc., Natick, MA) code developed in-house using a slicing thickness of 5 µm. Full-screen images were projected onto the vat window for photopolymerization. The printed layer thickness was 5 µm, and a printing time of 0.28 s was used for each layer. After printing, the parts were rinsed in ethanol to remove the unpolymerized ink and moved into a heating oven for thermal curing at 120 °C for 12 h.

#### Fabrication of DE-BVS

A MediCoat ultrasonic spray coating system (DES1000, Sono-Tek Corporation, Milton, NY) was used for coating the bare BVS with mPOC polymer. To formulate the spray coating ink, 8 wt.% mPOC pre-polymer was mixed with 0.55 wt% Irgacure 819 as the photoinitiator and 0, 1, 1.5, 2 and 4 wt% everolimus in the ethanol as solvent. The flow rate of 0.04 mL/min, ultrasonic power of 1.3 W, focusing gas pressure of 1 psi, distance of 5 mm between the nozzle and BVS, the rotation speed of 240 rpm, the horizontal translation speed of 0.1 inch per second, and drying gas pressure of 2 psi were optimized for obtaining a defect-free smooth coating on the surfaces of BVSs. After coating, BVSs were exposed to 365 nm UV light at 330 mW/cm^2^ for 2 min to cure the coating layers.

#### Scaffold characterization

The structure and surface morphology of the BVSs were observed by scanning electron microscope (SEM; Hitachi S4800, Japan). Strut thickness was measured from cross-sectional SEM images. The radial forces of the BVSs at 50% compression stain were measured by a radial force tester (RX650, Machine Solutions Inc., AZ). Fourier transformed infrared spectroscopy (FTIR) spectra of BVS coatings with and without drugs were acquired in the attenuated total reflection (ATR) model using a FTIR microscope (LUMOS by Bruker, Billerica, MA).

#### Drug release

DE-BVS and BVS (control) were immersed in the media of PBS containing 10% (v/v) ethanol in order to create an infinite sink condition for released everolimus at 37 °C under 150 rpm agitation. At scheduled time points, i.e. 6, 12, 24, 72 hours, 1, 2, 3, 4, 6, 8, 10, 12 weeks, the release media was collected and refreshed. The absorbance of collected media at 278 nm was measured using a microplate reader Cytation 5 (BioTek Instruments, Winooski, VT) according to the pre-determined standard curve.

#### Cell culture

Human umbilical vein endothelial cells (HUVECs; ATCC, Manassas, VA) were expanded in growth media consisting of Endothelial Cell Growth Kit-VEGF (ATCC PCS-100-041) in Vascular Cell Basal Medium (ATCC PCS-100-030) under the standard culture condition (37 °C with 5% CO2 in a humid environment) to 80% confluency before passaging. HUVECs at passages 5–7 were used. Human aortic smooth muscle cells (HAoSMCs; Cell Applications, Inc., San Diego, CA) were expanded in human SMC growth medium kit (Cell Applications, Inc., 311K-500) under the standard culture condition (37 °C with 5% CO2 in a humid environment) to 80% confluency before passaging.

#### Cell viability measurement

HUVECs or HAoSMCs were seeded onto 3D-printed substrates without (control) and with the coatings of drug or drug + barrier. After incubation for 24 h, cells were incubated with growth media containing alamarBlue cell viability reagents (Thermo Fisher Scientific, San Jose, CA) for 3 hours. After transferring reagents into 96-well plate, the fluorescence of each well was measured at 560/590 nm (excitation/emission) using a microplate reader Cytation 5 (BioTek Instruments). The number of viable cells per well was calculated against a standard curve prepared by plating various concentrations of isolated cells, as determined by the hemocytometer, in triplicate in the culture plates.

#### Immunostaining and imaging of cell morphology

Cells on flat and micropillar surfaces were fixed with 4% formaldehyde (Fisher Scientific, Fair Lawn, NJ) for 10 min and permeated with 0.1% Triton X-100 for 15 min. Then, the cells were incubated with Alexa Fluor® 594 Phalloidin (10 μM; A12381; ThermoFisher Scientific, Fair Lawn, NJ) and SYTOX™ Green Nucleic Acid Stain (0.15 μM; S7020; ThermoFisher Scientific) in PBS solution with 1.5% bovine serum albumin for 30 min. The samples were finally washed three times in PBS and mounted on microscope slides for examination using Leica spinning disk <u>confocal microscope</u>.

#### Animal model and implantation

Stent implantations were performed at North American Science Associates, LLC (NAMSA) (Minneapolis, MN), accredited by the Association for Assessment and Accreditation of Laboratory Animal Care International. Eight healthy farm swine (castrated male, 3-4 months, weight: 44-54 Kg) underwent device implantation with a targeted balloon-to-artery ratio of 1.0-1.1. Each animal received a single BVS, DE-BVS, or XIENCE DES (3.0 x 8 mm) for 28 days in 3 main coronary arteries, including left anterior descending coronary artery (LAD), left circumflex artery (LCX), and right coronary artery (RCA). RCA of one swine could not be accessed during interventional procedures, and thereby 1 of 8 XIENCE DESs could not be deployed. One of 8 DE-BVS was retrieved following the deployment in LAD of another swine due to the scaffold migration. A total of 22 devices (8 BVSs, 7 DE-BVSs and 7 XIENCE DESs) were implanted in 8 swine for 28 days. Intravascular ultrasound (IVUS) was performed before and after the procedure of device implantation and repeated after implantation for 14 and 28 days. 8 swine were euthanized and hearts were harvested for analysis.

#### Histological preparation and staining

Hearts were excised and perfusion fixed with 10% neutral buffered formalin at 80-100 mm Hg. The implanted coronary arteries were dissected from the heart, radiographed, and dehydrated in a graded series of ethanol solutions. Each artery was trimmed into three pieces, proximal native section, artery sections with the scaffold/stent, and distal native section, with the proximal and distal native sections aiming to be 10 mm away from the ends of the scaffold/stent. All artery sections were embedded in paraffin (all native arteries, 6 arteries with BVS, 5 arteries with DE-BVS) or plastic (all 7 arteries with XIENCE DES, 2 arteries with BVS, 2 arteries with DE-BVS). Then, 2-3 mm sections of arteries with scaffold/stent were cut, and 3 segments from the proximal, middle, and distal portions of each scaffold/stent were sectioned. Paraffin or plastic-embedded tissue sections were cut to 5 μm thickness. One of 7 arteries with XIENCE DES could not be cut into thin tissue sections due to the inappropriate plastic embedding. A total of 8 BVSs, 7 DE-BVSs, and 6 XIENCE DESs were sectioned for histological and immunohistochemistry (IHC) staining. Histomorphometry analysis was performed by staining tissues with hematoxylin and eosin (H&E) and Movat Pentachrome and light microscopic imaging. Inflammation was evaluated by IHC staining of two markers of macrophage – CD 86 and CD163. Vascular tissue recovery was evaluated by IHC staining of alpha-smooth muscle actin (α-SMA) of smooth muscle cells and VE-cadherin of endothelial cells.

#### Histomorphometry analysis

Histological sections were measured with computer assisted morphometric analysis by Fiji ImageJ software (https://imagej.net/Fiji), including EEL area (area bound by EEL), IEL area (area bound by IEL), luminal area (area bound by luminal border). After measurements were captured, the following calculations were made, when possible: intimal area (IEL area – luminal area), intimal thickness ({square root(IEL area/pi)} – {square root(luminal area/pi)}), and area stenosis ({1-(luminal area/IEL area)}x100). Inflammation score was calculated by the experienced pathologist based on the following grading parameters:

0 = no visible inflammatory response;

1 = Individual cells to isolated clusters of cells at the intima/subintima;

2 = Individual cells to isolated clusters of cells infiltrating the tunica media;

3 = increased numbers of cells/clusters with deeper tunica media infiltration;

4 = sheets of cells infiltrating the tunica media with effacement of wall features;

#### Statistics and reproducibility

Unless otherwise specified, data was presented as mean ± standard deviation (s.d.). For each experiment, at least three samples were analyzed. One-way ANOVA with Tukey’s Multiple Comparison Test was used to analyze statistical significance. A *P* value of <0.05 was considered to indicate a statistically significant difference. All experiments presented in the manuscript were repeated at least as two independent experiments with replicates to confirm the results are reproducible.

