## Supporting Information for "3D-printed, Citrate-based Bioresorbable Vascular Scaffolds for Coronary Artery Angioplasty"

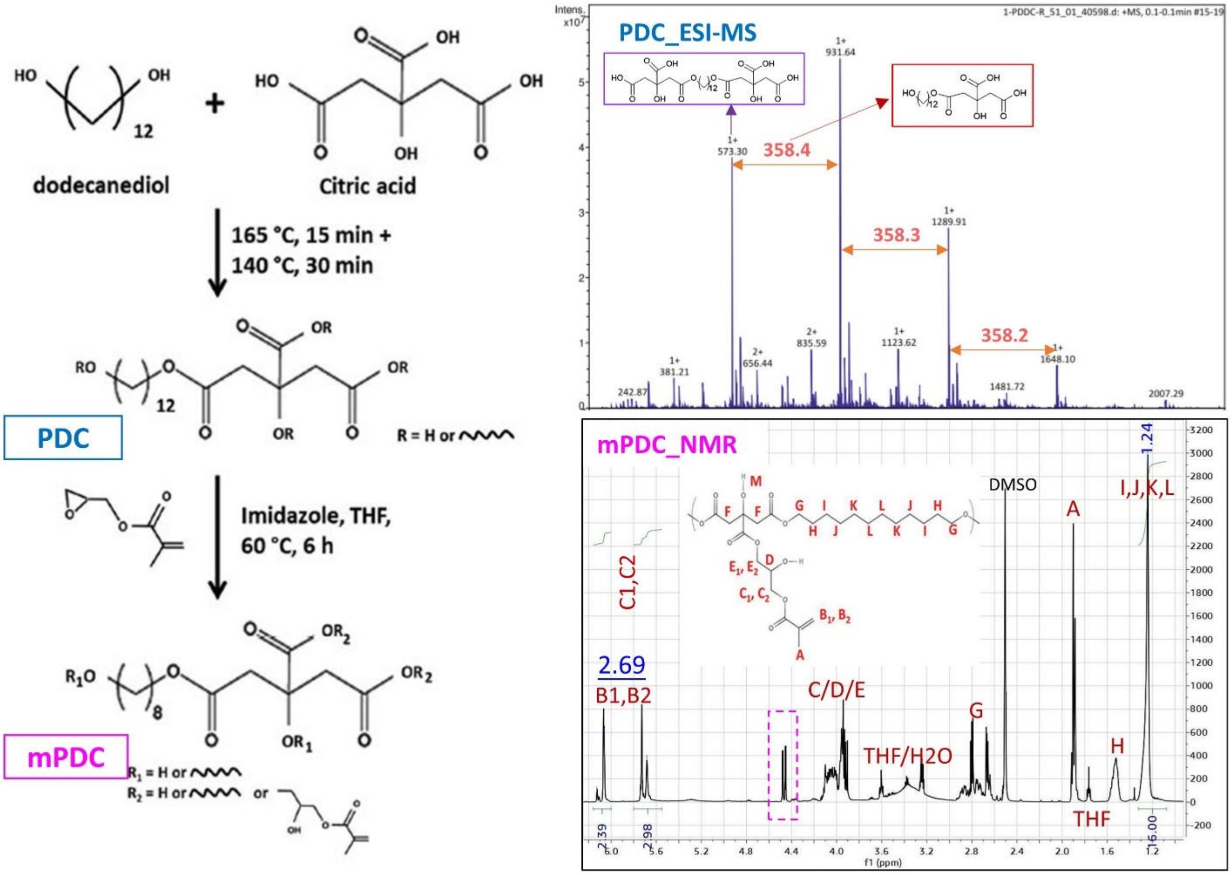


**Figure S1. Chemical synthesis and characterization of Methacrylated poly(1,12 dodecamethylene citrate) (mPDC).**

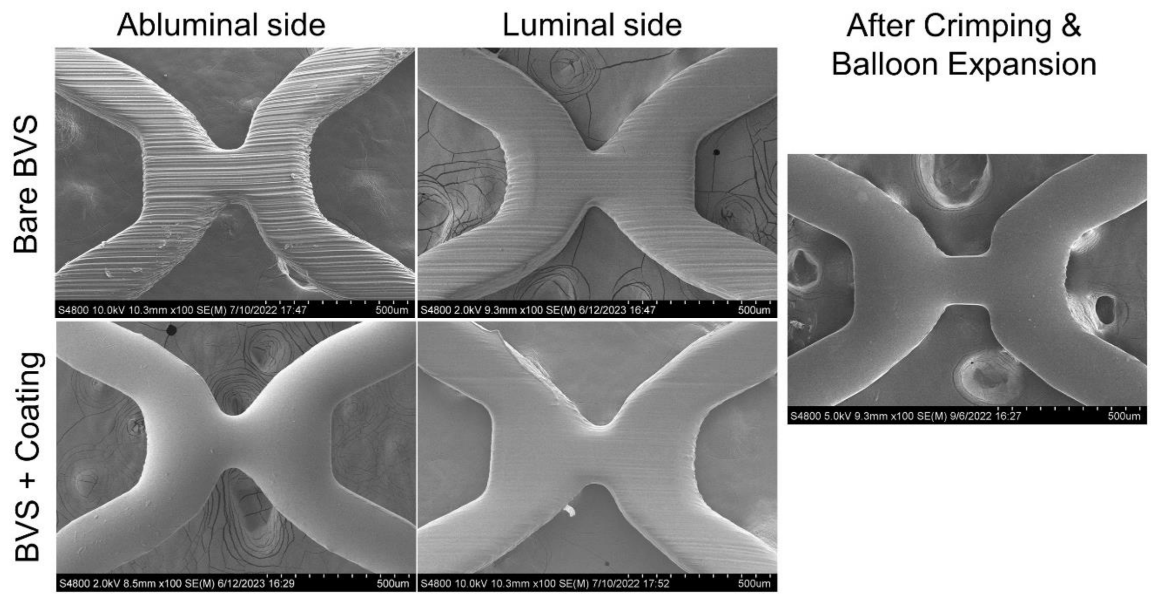


**Figure S2. SEM images show the deposition of smooth and uniform mPOC coatings on both luminal and abluminal sides of BVS.**

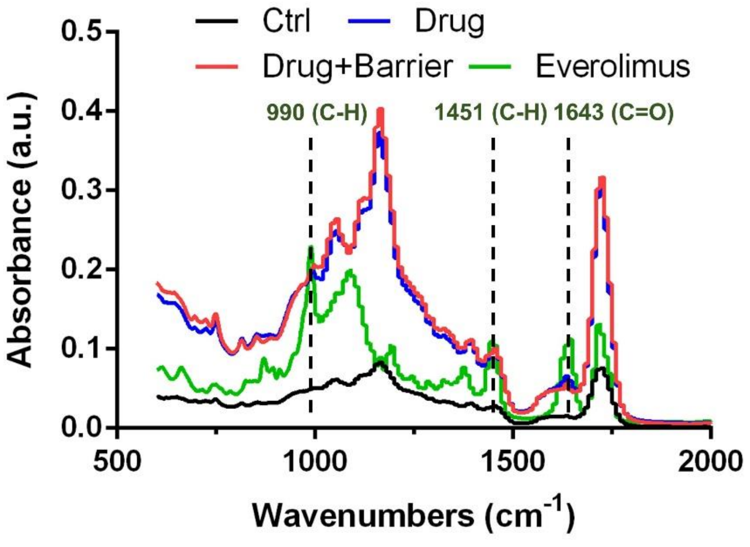


**Figure S3. FTIR spectra of pure everolimus and the BVS coatings without (Ctrl) and with drugs (Drug and Drug+Barrier).** The characteristic peaks of the everolimus were marked.

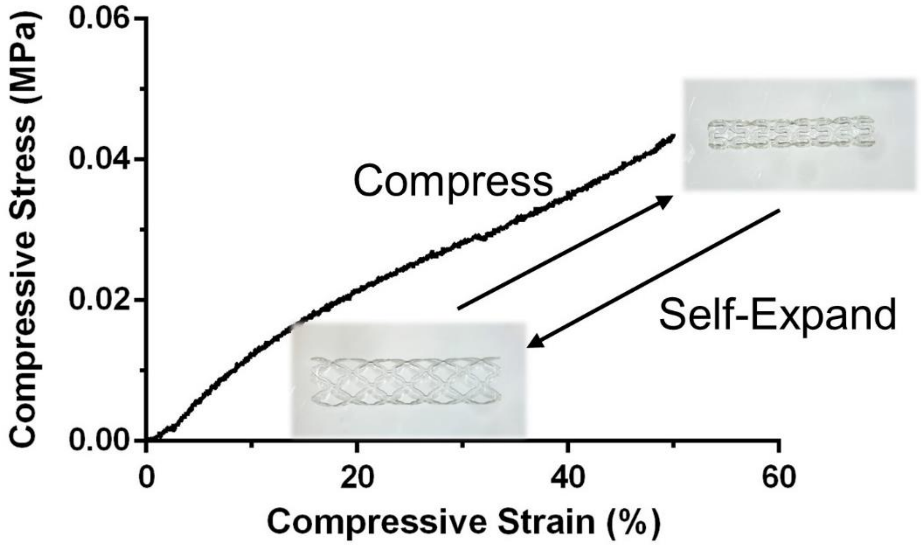


**Figure S4. The measured compressive stress-stain curve during the BVS compression process and the photos of compressed and self-expanded BVS.**

|  | IEL Area (mm²) |  |  | Lumen Area (mm²) |  |  | Neointimal Area (mm²) |  |  | Neointimal Thickness (mm) |  |  | Area Stenosis (%) |  |  | Inflammation Score |  |  |
|---|---|---|---|---|---|---|---|---|---|---|---|---|---|---|---|---|---|---|
| **Pig #** | **BVS** | **DE-BVS** | **XIENCE** | **BVS** | **DE-BVS** | **XIENCE** | **BVS** | **DE-BVS** | **XIENCE** | **BVS** | **DE-BVS** | **XIENCE** | **BVS** | **DE-BVS** | **XIENCE** | **BVS** | **DE-BVS** | **XIENCE** |
| **P1** | 4.87 | 2.80 | NA | 0.83 | 1.37 | NA | 4.04 | 1.43 | NA | 0.75 | 0.28 | NA | 83.00 | 51.02 | NA | 2.00 | 0.50 | NA |
| **P2** | 5.19 | 4.62 | NA | 1.69 | 1.54 | NA | 3.50 | 3.08 | NA | 0.55 | 0.54 | NA | 67.40 | 66.60 | NA | 2.33 | 3.00 | NA |
| **P3** | 5.85 | 5.27 | 4.11 | 2.32 | 2.82 | 2.38 | 3.53 | 2.45 | 1.73 | 0.51 | 0.35 | 0.27 | 60.36 | 46.53 | 42.09 | 3.00 | 2.00 | 3.00 |
| **P4** | 8.60 | 10.96 | 5.67 | 1.44 | 3.51 | 1.86 | 7.16 | 7.45 | 3.81 | 0.98 | 0.82 | 0.57 | 83.30 | 67.98 | 67.25 | 4.00 | 4.00 | 4.00 |
| **P5** | 4.41 | 4.09 | 4.92 | 2.79 | 2.25 | 3.61 | 1.62 | 1.84 | 1.31 | 0.24 | 0.30 | 0.18 | 36.72 | 44.97 | 26.62 | 2.00 | 2.00 | 1.00 |
| **P6** | 7.61 | 14.23 | 5.17 | 0.49 | 0.64 | 0.84 | 7.12 | 13.59 | 4.33 | 1.16 | 1.68 | 0.77 | 93.56 | 95.47 | 83.83 | 2.00 | 2.00 | 2.00 |
| **P7** | 4.98 | 4.06 | 6.11 | 3.14 | 1.62 | 3.87 | 1.84 | 2.44 | 2.24 | 0.26 | 0.43 | 0.29 | 36.95 | 60.16 | 36.66 | 2.00 | 2.67 | 2.67 |
| **P8** | 5.56 | NA | 6.00 | 3.65 | NA | 3.89 | 1.91 | NA | 2.11 | 0.25 | NA | 0.27 | 34.38 | NA | 35.19 | 0.67 | NA | 2.00 |

**Table S1. Histopathologic measurements at day 28**. Each parameter was measured and calculated by the mean of the proximal, middle, and distal segments of stent arteries. NA: data is not available (see details in the Methods and Materials).
